# Intentional gestures predict complex sociality in wild chimpanzee

**DOI:** 10.1101/365858

**Authors:** Anna Ilona Roberts, Sam George Bradley Roberts

**Author notes:** Equally contributing authors.

## Abstract

A key challenge for primates is coordinating behavior with conspecifics in large, complex social groups. Gestures play a key role in this process and chimpanzees show considerable flexibility communicating through single gestures, sequences of gestures interspersed with periods of response waiting (persistence) and rapid sequences where gestures are made in quick succession, too rapid for the response waiting to have occurred. Previous studies examined behavioral reactions to single gestures and sequences, but whether this complexity is associated with more complex sociality at the level of the dyad partner and the group as a whole is not well understood. We used social network analysis to examine how the production of single gestures and sequences of gestures was related to the duration of time spent in proximity and individual differences in proximity in wild East African chimpanzees (*Pan troglodytes schweinfurthii*). Pairs of chimpanzees that spent a longer duration of time in proximity had higher rates of persistence, but not a higher rate of single gesture or rapid sequences. Central individuals in the social network received higher rates of persistence, but not rapid sequence or single gesture. Intentional gestural communication plays an important role in regulating social interactions in complex primate societies.

## Introduction

Primate social life has frequently been described as particularly complex in its nature and when compared with other vertebrates, primates have unusually large brains for their body size (Dunbar 1993; Dunbar 1998). Primate sociality is based on bonded social relationships where individuals repeatedly interact with the same group members in many different contexts (Freeberg et al. 2012). It has been proposed that the sociality of primates is cognitively demanding, leading to evolution of large brains in both primates and hominins (Dunbar and Shultz 2007a). In particular, there is a strong positive correlation between group size and brain size in primates, and particularly neocortex size in relation to the rest of the brain (Dunbar 1993). Thus, primates living in larger groups have larger neorcortex ratios (Dunbar and Shultz 2007a). The relationship between brain size and group size may be influenced by the demands arising from maintaining social relationships in primates. Primates use grooming behavior to maintain stable, long lasting, and differentiated social relationships with both related and unrelated individuals (Dunbar 2010). The time and cognitive demands arising from maintaining social relationships through grooming result in a multilevel group structure, with hierarchically nested layers of social bonds, delineated by decreasing amounts of time spent in grooming behaviour (Hill et al. 2008).

In addition, gestural communication, defined as voluntary movements of the arms, head, body postures and locomotory gaits (Bard 1992; Hewes 1973; Roberts et al. 2014a; Tomasello et al. 1984) is important in maintaining social relationships of primates (Bard 1992; Bard et al. 2014; Forrester 2008; Fröhlich et al. 2016; Genty et al. 2009; Gillespie-Lynch et al. 2013; Halina et al. 2013; Hewes 1973; Hobaiter and Byrne 2011a; Leavens et al. 2005; Liebal et al. 2004; Maestripieri 2005; McCarthy et al. 2012; Pika et al. 2005; Pollick and de Waal 2007; Roberts et al. 2014a; Roberts et al. 2012b; Schneider et al. 2012; Scott 2013; Taglialatela et al. 2015; Tomasello et al. 1984; Tomasello et al. 1985). Gestural communication is particularly relevant for studies of social cognition because gestures can influence social bonding through intentional behaviour or emotional expression and this may have important implications for the complexity of cognitive skills involved in managing of social relationship. In intentional gesturing, signallers have a goal and influence the recipient flexibly based on an understanding that recipients have goal states different from their own and these states can affect their behaviour (Tomasello and Zuberbühler 2002). In addition, gestures can coordinate social bonding behaviour by fulfilling social bonding function in itself and simply expressing the signaller’s affect. These emotional gestures may not be contingent upon the signaller’s goal but are diffuse expressions of signaller’s internal emotional state that can release social bonding neurohormones in the recipients (Dunbar 2010). For instance, greeting gestures when encountering each other after a period of separation can influence social bonding with the recipient and hence influence duration of time spent in close proximity. Thus, emotional communication has an adaptive function and can coordinate social behaviour because it influences emotional states of the recipients (Spoor and Kelly 2004).

In particular, primate gestures that occur singly or in sequences can reveal the link between gestural communication and social bonding (Cartmill and Byrne 2007a; Genty and Byrne 2009; Hobaiter and Byrne 2011b; Leavens et al. 2005; Liebal et al. 2004; McCarthy et al. 2012; Roberts et al. 2014a; Roberts et al. 2012a; Roberts et al. 2013; Roberts et al. 2014b; Tanner 2004; Tanner and Perlman 2016; Tempelmann and Liebal 2012; Tomasello et al. 1984) Series of gestures made in anticipation of a response, as shown by persistence (Gómez 1996; Moore 2016; Scott-Phillips 2015a; Scott-Phillips 2015b) may be important in social bonding in primates because they are made intentionally (Cartmill and Byrne 2007a; Leavens et al. 2005; Roberts et al. 2013; Roberts et al. 2014b). In gestural communication that is characterized by persistence, the signaller makes a gesture, pauses for one to five seconds to wait for a response, and then if the response is not forthcoming, the signaller makes another gesture (Hobaiter and Byrne 2011b). Moreover, great apes can also make a ‘rapid sequence’ whereby several gestures are made in quick succession, too rapid for the response waiting to have taken place (Hobaiter and Byrne 2011b).

In intentional communication the signaler modifies the production of the signals flexibly (Bates et al. 1979; Leavens et al. 2005; Tomasello et al. 1984). In support of this role of gestures, observational and experimental research in experimental tasks, and in conspecific social interactions, showed that signalers can adjust their gestural communication in relation to the changes in the behaviour of the recipient (Cartmill and Byrne 2007a; Genty and Byrne 2009; Hobaiter and Byrne 2011b; Leavens et al. 2005; Liebal et al. 2004; McCarthy et al. 2012; Roberts et al. 2014a; Roberts et al. 2012a; Roberts et al. 2013; Roberts et al. 2014b; Tanner 2004; Tanner and Perlman 2016; Tempelmann and Liebal 2012; Tomasello et al. 1984). In experimental studies that manipulated the response consequences of ‘unsuccessful’ communication against a baseline of ‘successful’ communication, it was clearly demonstrated that apes can respond to the different behavioural states of the experimenter (Cartmill and Byrne 2007b; Leavens et al. 2005). For instance, individuals discontinued communicative attempts when the desired response was obtained and continued communicating when faced with an absence of a response (Cartmill and Byrne 2007a; Cartmill and Byrne 2010; Leavens et al. 2005; Roberts et al. 2012a; Roberts et al. 2013; Roberts et al. 2014b). Moreover, in a food finding task that required language-trained chimpanzees to guide a naïve human experimenter to a hidden food item, the chimpanzees coordinated their behavior with the experimenter in a flexible way, based on the experimenter’s responses to the chimpanzees’ communication. The chimpanzees used non-indicative gestures such as bobbing when the experimenter accurately pointed to the food location and indicative gestures such as pointing when the experimenter pointed to a location where the food was not hidden (Roberts et al. 2014b). However, whilst the role of persistence in influencing the recipient’s behaviour has been shown in previous studies, the role of persistence in social bonding is currently unclear. In addition, very little is known about the role of single gestures and rapid sequences in social bonding. Thus, the issue of whether great apes can use gestural communication flexibly to coordinate social behaviour with different types of social partners, and how this use relates to variations in social network size, remains unresolved.

Chimpanzees are an ideal species to examine the relationship between sociality and the production of single gestures, persistence and rapid sequences in primates. Chimpanzees live in complex fission-fusion groups, where association dynamics are fluid and chimpanzees form temporary subgroups (‘parties’) that vary in size, composition and duration (Goodall 1986). Due to this fission-fusion structure, patterns of interaction between pairs of chimpanzees can vary on daily basis. In this study we examine the relationship between social interactions and the production of single gestures, persistence and rapid sequences in wild East African chimpanzees (*Pan troglodytes schweinfurthii*) in Budongo Forest, Uganda using Social Network Analysis (SNA). We examine how different types of communication (single gesture, rapid and persistence sequence) are related to sociality. In this study, consistent with previous research in this area (Lehmann et al. 2016; Sapolsky et al. 1997; Silk et al. 2013; Silk et al. 2010b), we used proximity to measure differences in sociality between pairs of chimpanzees. We examined how these differences in sociality relate to patterns of communication between pairs of chimpanzees.

Through emotional communication signaler induces compatible affect in the recipient and through synchronized affect, the emotional communication facilitates attentional and behavioral convergence of the dyad partners (Owren and Rendall 2001). In contrast, intentional communication influences behavior of the recipient by influencing their movement and attention to achieve a goal such as travel to the same location. It has been argued that intentional communication has evolved as a means to enable social bonding with dyad partners as it can influence behavior of the recipient more flexibly than emotional gesture and this may have been accompanied by increase in brain size during the course of hominin evolution. In this study we explored the associations between proximity and different types of gestural communication. We hypothesize that proximity will be differentially associated with the rates of different types of gestural communication – single gestures, rapid sequences and persistent sequences. Specifically we predict that that intentional communication (e.g. single gesture, persistence sequence) will be associated with a longer duration of time spent in proximity relative to emotional gestures (e.g. rapid sequence) (Hypothesis 1).

However, it is unclear whether single gestures, rapid and persistence sequences differ in response types made to the gestures and this would indicate the degree to which these communication types are intentional. Recipients can respond in a goal directed way by adjusting behaviour to the goal conveyed in the gesture, but can also respond communicatively. Thus, we hypothesize that goal directed and communicative responses will be differentially associated with the type of communication (Hypothesis 2). We predict that intentional gestures (single gesture, persistence) will be associated with goal directed response (by activity change) whereas emotional gestures (rapid sequence) will be associated with response by communication (visual, tactile gesture or vocalisation).

Furthermore, it is currently unclear whether the response to the gesture may be associated with the degree of sociality. Presence and type of response (e.g. goal directed or communicative) can indicate the willingness of the recipient to coordinate behaviour with the signaller and thus reflect the level of social bonding (Schneider et al. 2017; Wilke et al. 2017). Following on from Hypothesis 1, we hypothesize that the presence (Hypothesis 3) and type (Hypothesis 4) of response will be associated with sociality. Specifically, we predict that if intentional gestures facilitate social bonding then we would expect longer duration of time spent in proximity to be associated with higher rate of response present and response by activity change. In contrast shorter duration of time spent in proximity would be associated with higher rate of response absence and response by communication.

Finally, individuals have different positions in the group, with central individuals having more social bonds relative to peripheral individuals who have fewer social bonds (Roberts and Roberts 2016a; Roberts and Roberts 2016b). Previous research has suggested that more central individuals have different overall patterns of communication to peripheral individuals (Roberts and Roberts 2016a; Roberts and Roberts 2016b). We therefore predict that the centrality of individual chimpanzees will be associated with the rate of singe, rapid and persistent gestural communication they produce and they receive (Hypothesis 5).

The relationship between communication and social behaviour could arise simply as a relation between a behaviour that requires proximity with a metric of proximity. To avoid this possibility, in all analyses we control for the duration of time spent in close proximity (all communication indices are calculated per duration of time spent within 10 m). Furthermore, in addition to the sequence type, biological factors such as reproductive status, age similarity, sex similarity and kinship have been shown to influence patterns of social bonding between pairs of chimpanzees (Langergraber et al. 2009; Mitani 2009; Roberts and Roberts 2016b). Thus we control for these biological factors in all models.

## Methods

### Study site and subjects

The behaviour of East African chimpanzees (*Pan troglodytes schweinfurthii*) of the Sonso community at the Budongo Conservation Field Station, Budongo Forest Reserve in Uganda (latitude 1° 37’-2° 00’N; longitude: 31° 22’-31°46’E) was observed in relation to communication and social relationships between March and June 2008, following subjects between 07:00 and 16:00 at least 5 days a week. The distance to the focal chimpanzee and the limb injuries of the chimpanzee can influence the frequency and type of gestural communication. Thus from the community of approximately 74 individuals including 21 adult females and 10 adult males, a sample group of 12 adult focal subjects (6 adult males and 6 adult females) was chosen to ensure lack of any limb injuries and in accordance with the level of habituation, simultaneously ensuring that age and rank classes were equally represented in the sample – see Table 1 (Roberts and Roberts 2016b) for demographic and sampling details of the focal chimpanzees. The study was non-invasive and the study methods were approved by the University of Stirling Ethics Committee. Full details of the study site, subjects, data collection, video analysis and classification of gestures have been described previously (Roberts et al. 2014a), so only the key information is provided here.

**Table 1.** Focal ID, sex, year of birth and reproductive status of the 12 focal subjects included in the study.

| **Focal subject ID** | **Sex** | **Age** | **Female reproductive status** | **Total observation duration (minutes)** |
| --- | --- | --- | --- | --- |
| BB | Male | 21 | - | 516 |
| HW | Male | 15 | - | 1030 |
| KT | Male | 15 | - | 1026 |
| KU | Female | 29 | Pregnant | 910 |
| KW | Female | 27 | Nursing | 510 |
| ML | Female | 33 | Cycling | 1118 |
| MS | Male | 17 | - | 524 |
| NB<sup>c</sup> | Female | 46 | Cycling | 500 |
| NK<sup>a</sup> | Male | 26 | - | 582 |
| RH | Female | 43 | Nursing | 1038 |
| SQ | Male | 17 | - | 554 |
| ZM | Female | 40 | Cycling | 710 |

Notes. <sup>a</sup> Alpha male' <sup>b</sup> Alpha female.
Dominance based on unidirectional pant-grunt calls – for full details, see (Roberts and
Roberts 2016b)

### Data collection protocol

During 18-minute focal follows consisting of 9 scans (nine 2-minute intervals), two types of social information were recorded. First, the association and activity patterns were recorded. These included the identity of individuals present within 10 m and more than 10 m away from the focal individual, and the identity, visual attention, distance and activity of the nearest neighbour to the focal individual. Second, gestural communication to accompany the 18-minute instantaneous sampling of association and behaviour patterns in the chimpanzees was recorded continuously using a digital video camera recorder.

Visual attention between the focal individual and the nearest neighbour was recorded using categories presented in Supplementary Information 2. We tested the similarity in association patterns between the scans taken at 2 minute intervals, to examine the extent to which association patterns changed during the 18 minute focal follows, and between one focal follow and the next. For full details of this analysis, see (Roberts and Roberts 2016a; Roberts and Roberts 2016b). Briefly, the results demonstrated that the adjacent scans taken at 2 and 4 minutes of the 18-minute sampling period yielded similar findings, and thus adjacent 2 minute scans within a focal follow were treated as continuous data. However, the first scan (2 min) and final scan (18 min) during the focal follow differed both for 10 m associations and party level associations. Thus the association patterns change significantly over the course of an 18-minute focal follow, meaning each 18-minute focal follow can be considered an independent sample of association patterns.

### Behavioural measures

First, we used the genetic relationships identified in previous studies to classify pairs (dyads) of chimpanzees as kin or non-kin (Reynolds 2005), taking into account maternal kin relations only (relatedness 0.5). We classified dyads of chimpanzees as belonging to the same (5 years or less age difference) or a different (above 5 years age difference) age class (Mitani et al. 2002) and also according to reproductive and sex similarity. The details of the categorization of attribute data are provided in Table 2.

**Table 2.** Variables included in the models

| Independent variable | Definition | Frequencies or mean±SD/ 95% CI<br>(duration/frequency per hour spent within 10 meters) |
| --- | --- | --- |
| Persistence sequence | A series of gestures whereby there are pauses of 1 -5 seconds between consecutive gestures | 0.11±0.45, [0.03, 0.18] |
| Single gesture | A single gesture that is not made in series and where there is at least 30 seconds to the next consecutive gesture | 1.27±4.07, [0.57, 1.97] |
| Rapid sequence | A series of gestures without pauses between consecutive gestures | 0.45±1.30,<br>[0.23, 0.68] |
| Sex difference | Sex difference between focal subject and the recipient (0 = different sex: male-female or female-male, 1 = same sex: male-male or female-female) | 0 = 60, 1 = 60 |
| Age difference | Age difference between focal subject and the recipient (0 = different age: more than 5 years age difference between individuals in the dyad, 1 = same age: no more than 5 years age difference between individuals in the dyad) | 0 = 102, 1 = 30 |
| Oestrous similarity | Reproductive state difference between focal subject and the recipient (0 = reproductively inactive: unoestrous female- unoestrous female, unoestrous female-oestrous female, oestrous female-oestrous female, unoestrous female-male, male-male; 1 = reproductively active: male-oestrous female) | 0 = 96, 1 = 36 |
| Maternal kinship | Maternal kinship presence between focal subject and the recipient (0 = unrelated dyad, 1 = mother-son; son-mother) | 0 = 126, 1 = 6 |
| Proximity | Duration of time individual spent in proximity within 10 metres per hour spent in the same party | 23.26±1.22,<br>[20.84, 25.69] |
| Response by activity change | Change of behaviour by means of goal directed response, whereby recipient performs some action that conforms to the goal of the signaller (e.g. starts to groom) | 0.58±1.80,<br>[0.26, 0.89] |
| Response by vocalization | Change of behaviour by means of vocalization (production of sound via vocal tract) by the recipient, | 0.47±2.02,<br>[0.12, 0.82] |
|  | which is not followed by goal directed action towards signaller (e.g. pantgrunt during travel, whereby signallers travel before and after the pantgrunt) |  |
| Response by visual or tactile gesture | Change of behaviour by means of visual or tactile gesture which excludes production of sound by the recipient via vocal tract. This behaviour is not followed by goal directed action towards signaller (e.g. embrace during travel, whereby signallers travel before and after the embrace) | 0.08±0.40, [0.01, 0.14] |

Second, to establish the rates of gestures between dyads, the video footage was viewed on a television and the cases of nonverbal behaviour that were identified were coded as an act of gestural communication if they met following criteria: 1) the non-verbal behaviour was an expressive movement of the limbs or head and body posture that was mechanically ineffective, 2) the behaviour was communicative by non-mechanical means (i.e. consistently produced a change in the behaviour of recipient or facilitated maintenance of activity, e.g. grooming). Whilst the criterion of ‘non-mechanical means’ did not exclude cases of physical bodily movement by the signaller of a social partner, it was important that such cases had a communicative purpose, i.e. rather than just move the body part of the social partner physically, these cases also displayed communicative purpose, For example during grooming, the light touch of the body and subsequent slight displacement of the body part also meant the desire for the social partner to move the body part. Next, behaviour had to be goal directed to be considered intentional (Bard 1992; Bates et al. 1979). The intentionality of gestures was coded sensu Tomasello et al. (Tomasello et al. 1985) who gave following example to explain intentionality of gestures: ‘a child might be struggling to open a cabinet, crying and whining as s/he struggles. Seeing this, the mother might come to the rescue and open the cabinet. This is a perlocutionary act because, while communication may be said to have occurred, the “sender” (the child) did not intentionally direct any behavior towards the mother. If, on the other hand, the child has turned its attention from the cabinet to the mother and whined at her, the whining now becomes a social-communicatory act with the intention of obtaining adult aid’. Operationally, thus, one clear evidence for intentionality of gestures comes from the presence of an audience and visual attention between signaller and the recipient during production of the gesture. In this dataset, all cases of gesturing included the presence of an audience in close proximity (Supplementary Information 1 and 2), so the intentionality of the gestures in this dataset was not differentiated by the presence of the audience. In addition, the presence and absence of bodily orientation before and during the gesture were coded to establish intentionality of gestures (see Supplementary Information 2 for details for each gesture type). The presence and absence of communicative persistence was also coded in this paper following communicative persistence sensu Hobaiter and Byrne (Hobaiter and Byrne 2011a; Townsend et al. 2016). In order to establish communicative persistence, gesture events were scored in accordance to whether they occurred singly or in sequences, defined as one or more than one gesture made consecutively by one individual, towards the same recipient, with the same goal, within the same context, and made within a maximum of 30 seconds interval to ensure independence. According to the classification by Hobaiter and Byrne (Hobaiter and Byrne 2011b), persistence of gesturing is when the chimpanzee produces one gesture or a gesture sequence, then after a period of response waiting (1-5s) they produce another gesture – here such instances are termed a ‘persistence sequence’. However, when a chimpanzee produces a sequence and there is no intermittent pause between gestures, then the chimpanzee has not persisted – here such instances are here termed a ‘rapid sequence’. Supplementary Information 2 contains detailed information for the percentages of each gesture type occurring within each sequence type. Moreover, Supplementary Information 1 (Table 2) provides the number of cases of single gestures, persistence and rapid sequences per each focal subject separately. The panthoot behaviour is broadcast at a wider audience and within social network analysis we counted all individuals present within 10 meters as recipients of any gestures accompanied by pant hoots produced by the focal subject. The identity of the recipients of the panthoot was taken from the scan sample recorded every 2 minutes.

A random sample of 50 sequences of gestures was coded by a second coder for intentionality (response waiting and persistence) and the Cohen’s Kappa coefficient showed good reliability (K = 0.74) (Bakeman and Gottman 1997). In this sample of reliability coding of persistence, one requirement for categorizing the event as persistence was the presence of mutual bodily orientation between the signaller and the recipient. Thus in this sample, response waiting and persistence co-occurred in all cases of gesturing.

Having established the independence of the data collection protocol, the behavioural measures for each dyad of the signaller and the recipient were calculated in the following manner:

### The dyadic communication measure

The dyadic communication measure (CA) is the rate at which focal subject A communicated to non-focal subject B when B was in close proximity (within 10 m) to focal subject A, per hour spent within 10 m of the non-focal subject B, or:

CA_AB_ = (C_AB_* 60) / P10_AB_ *2

where C_AB_ = the number of times A communicated with B when in close proximity (within 10m) to B

P10_AB_ = the number of times A was in close proximity (within 10m) to B

2 = duration of instantaneous subsample interval in minutes

60 = the number of minutes in an hour

Social Network Analysis (SNA)

The behavioral measures were entered into a network matrix consisting of 12 rows and 12 columns, with each row and column designating a different focal chimpanzee. In this analysis only data on 132 focal and non-focal subject dyads was included in the analysis, excluding any data where the recipient was not a focal subject in this study. The number of entries for each behavioural measure are provided in Table 2. The values in each cell of the matrix represented the value for communication or proximity for a specific pair of chimpanzees (e.g. the rate of persistence sequence between Bwoba and Hawa, per hour spent within 10m). These networks were weighted – i.e. each cell consisted of a continuous value representing that behaviour, rather than a 1 or a 0 indicating the presence or absence of a tie. Further, the networks were directed in that the rate of gestures by Bwoba that were directed to Hawa may be different from the rate of gestures by Hawa that were directed to Bwoba.

The observations that make up network data are not independent of each other and thus in general standard inferential statistics cannot be used on network data. Instead, a set of analyses using randomisation (or permutation) tests have been developed where the observed value is compared against a distribution of values generated by a large number of random permutations of the data. The proportion of random permutations in which a value as large (or as small) as the one observed is then calculated, and this provides the *p* value of the test (Borgatti et al. 2013). We used Multiple Regression Quadratic Assignment Procedure (MRQAP) to examine the relationships between the networks (Borgatti et al. 2013). MRQAP regression is similar to standard regression in that it allows for the examination of the effect of a number of independent variables (e.g. gestural communication network) on an outcome variable (e.g. proximity network). Several different types of MRQAP regression are available and we used Double Dekker Semi-Partialling MRQAP regression, which is more robust against the effects of network autocorrelation and skewness in the data (Dekker et al. 2007). The number of permutations used in this analysis was 2,000. All data transformations and analyses were carried out using UCINET 6 for Windows (Borgatti et al. 2014).

## Results

### Type of sequence

We examined a total of 545 sequences (1044 instances of gestures) performed by 12 focal adult individuals towards other focal and non-focal adult individuals to examine the extent to which the gestures presented in this dataset were intentional. The percentage of association between each gesture type separately and indices of intentionality is given in Supplementary Information 1, Table 1. Moreover, frequencies of gesture events within these categories are provided in Supplementary Information 2. In this sample (consisting of adult to adult gestures only) the mean percentage ± SD [95% CI] of cases of all gesture types associated with the presence of bodily orientation by the signaller towards the recipient during production of the gesture was 91.5 ± 18.5%, [87, 95]. The mean percentage ± SD [95% CI] of cases of all gesture types associated with the presence of recipients’ bodily orientation towards signaller, when the signaller’s bodily orientation towards the recipient was absent, was 6.9 ± 15.4% [3, 10]. Finally, the mean percentage ± SD [95% CI] of cases of all gesture types where neither signaller nor the recipient were bodily oriented towards one another during production of the gesture was 1.5 ± 11% [0, 3]. This shows that the gestures in our dataset were communicative and intentional according to the previously established criteria for defining intentionality in preverbal humans and primates (Bard 1992; Bates et al. 1979). In this paper, sequences were categorized as either single gestures, persistence sequences or rapid sequences following Hobaiter and Byrne (Hobaiter and Byrne 2011b), taking into account both manual and bodily gestures (Roberts et al. 2014a; Roberts et al. 2012b). Per focal individual, the mean number ± SD [95% CI] of single gestures was 32.0 ± 32, [11.69, 52.47], for persistence sequences was 4.41 ± 5.85, [0.69, 8.13] and for rapid sequences was 8.9 ± 9.09, [3.14, 14.69] – see also Supplementary Information 1, Table 2 for frequency of single gestures, persistence and rapid sequences for each focal subject separately. In this study we used two main sets of analyses: Multiple Regression Quadratic Assignment Procedures (MRQAP) and node-level regression. The description of all the variables included in these models are provided in Table 2. In all analyses, the age, sex, reproductive status, kinship were included in the models, including the recipient of the gesture entered as a dyad partner in all models. Only statistically significant findings are presented in this block of results. Full details of the models including all variables are provided in Tables 3 - 9.

**Table 3.** MRQAP regression models showing predictors of proximity (duration of time spent within 10 meters per hour spent in same party) by sequence type of gestures between N = 12, 132 dyadic relationships of the chimpanzees. Significant P values are indicated in bold.

| Attribute category/ rate of gesture sequence per hour spent in close proximity | Standardized coefficient | Standard error | P |
| --- | --- | --- | --- |
| Age similarity | 0.162 | 3.658 | 0.060 |
| Sex similarity | -0.091 | 3.760 | 0.239 |
| Kinship similarity | 0.065 | 6.742 | 0.258 |
| Oestrous similarity | 0.006 | 4.328 | 0.487 |
| Rapid sequence | -0.025 | 1.107 | 0.389 |
| Single gesture | 0.110 | 0.370 | 0.138 |
| Persistence sequence | 0.164 | 3.109 | **0.034** |

**Table 4.** MRQAP regression models showing predictors of rapid sequence (rate of production per hour spent within 10 meters) by rate of response to the gesture between N = 12, 132 dyadic relationships of the chimpanzees. Significant P values are indicated in bold.

| Attribute category/ rate of gesture<br>sequence per hour spent in close<br>proximity | Standardized<br>coefficient | Standard error | P |
| --- | --- | --- | --- |
| Age similarity | 0.010 | 0.160 | 0.386 |
| Sex similarity | -0.057 | 0.169 | 0.176 |
| Kinship similarity | -0.037 | 0.283 | 0.142 |
| Oestrous similarity | -0.060 | 0.193 | 0.171 |
| Response by visual or tactile gesture | 0.006 | 0.353 | 0.471 |
| Response by activity change | -0.067 | 0.084 | 0.271 |
| Response by vocalisation | 0.857 | 0.065 | **0.001** |

**Table 5.** MRQAP regression models showing predictors of persistence sequence (rate of production per hour spent within 10 meters) by rate of response to the gesture between N = 12, 132 dyadic relationships of the chimpanzees. Significant P values are indicated in bold.

| Attribute category/ rate of gesture<br>sequence per hour spent in close<br>proximity | Standardized<br>coefficient | Standard error | P |
| --- | --- | --- | --- |
| Age similarity | -0.029 | 0.086 | 0.373 |
| Sex similarity | 0.042 | 0.086 | 0.327 |
| Kinship similarity | -0.015 | 0.152 | 0.437 |
| Oestrous similarity | 0.053 | 0.095 | 0.275 |
| Response by visual or tactile gesture | -0.754 | 0.181 | **0.001** |
| Response by activity change | 1.132 | 0.048 | **0.001** |
| Response by vocalisation | 0.067 | 0.019 | 0.134 |

**Table 6.** MRQAP regression models showing predictors of single gesture (rate of production per hour spent within 10 meters) by rate of response to the gesture between N = 12, 132 dyadic relationships of the chimpanzees. Significant P values are indicated in bold.

| Attribute category/ rate of gesture<br>sequence per hour spent in close<br>proximity | Standardized<br>coefficient | Standard error | P |
| --- | --- | --- | --- |
| Age similarity | 0.103 | 0.492 | **0.017** |
| Sex similarity | 0.047 | 0.493 | 0.195 |
| Kinship similarity | 0.002 | 0.844 | 0.373 |
| Oestrous similarity | 0.037 | 0.534 | 0.282 |
| Response by visual or tactile gesture | 0.392 | 0.901 | **0.001** |
| Response by activity change | 0.488 | 0.247 | **0.001** |
| Response by vocalisation | 0.068 | 0.100 | 0.083 |

### Type of sequence and proximity (Hypothesis 1)

We used MRQAP to examine the relationship between duration of time spent in proximity (within 10 meters per hour spent in same party) and the rate of production of gestures (frequency per hour spent within 10 m) and demography (Table 3). The proximity was significantly positively associated with a higher rate of persistence sequence between dyads (β = 0.164, *p* = 0.034). In contrast, the rate of rapid sequences or persistence sequences was not significantly associated with the proximity (Fig. 1).

**Fig 1.**
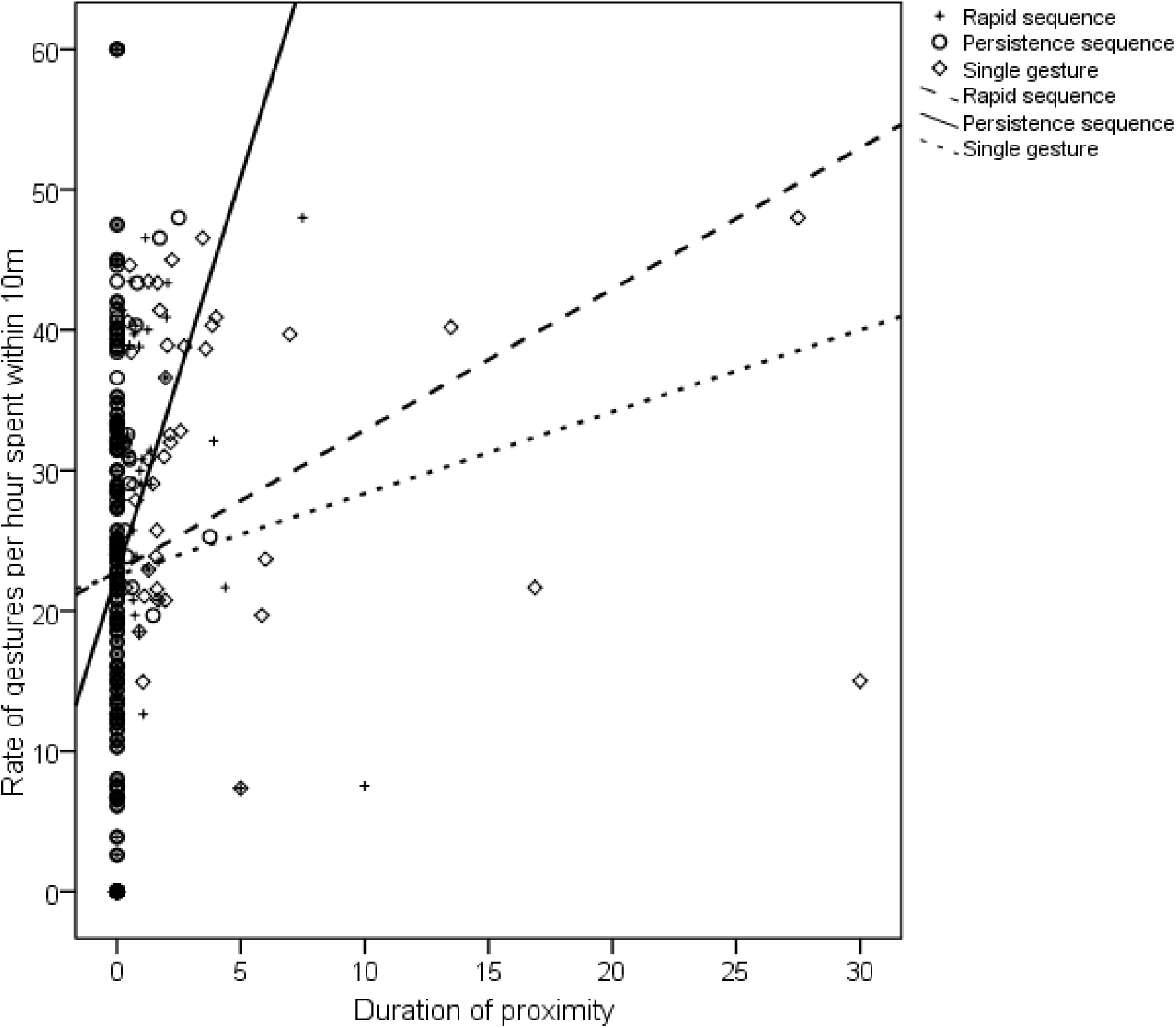
Proximity (duration of time spent within 10 meters per hour spent in the same party) and rate of single gestures, rapid sequences and persistence sequences in dyads of chimpanzees (n = 132).

### Type of sequence and type of response (Hypothesis 2)

We then examined how the rate of response type to the gestures (response by visual or tactile, gesture, response by vocalization, response by activity change) was associated with the type of sequence (rapid sequence, persistence sequence, single gesture) (Tables 4 - 6). There was a positive association between response by vocalization and rapid sequence (β = 0.857, *p* = 0.001). Moreover, there was a positive association between a single gesture and response type by activity change (β = 0.488, *p* = 0.001) and positive association between a single gesture and response by visual or tactile gesture (β = 0.392, *p* = 0.001). Finally, there was a positive association between response by activity change and persistence (β = 1.132, *p* = 0.001) but negative association between response by tactile or visual gesture and persistence (β = − 0.754, *p* = 0.001).

### Presence and absence of response and proximity (Hypothesis 3)

We next examined how the rate of response type to the gestures (response presence and absence) was associated with the proximity (Table 7) There was a significant positive association between proximity and response presence (β = 0.178, *p* = 0.026).

**Table 7.** MRQAP regression models showing predictors of proximity (duration spent within 10 meters per hour spent in same party) by rate of response present or absent to the gesture between N = 12, 132 dyadic relationships of the chimpanzees. Significant P values are indicated in bold.

| Attribute category/ rate of gesture sequence per hour spent in close proximity | Standardized coefficient | Standard error | P |
| --- | --- | --- | --- |
| Age similarity | 0.149 | 3.748 | 0.078 |
| Sex similarity | -0.059 | 3.704 | 0.321 |
| Kinship similarity | 0.064 | 6.619 | 0.252 |
| Oestrous similarity | 0.030 | 4.282 | 0.397 |
| Response absent | 0.006 | 0.573 | 0.466 |
| Response present | 0.178 | 0.380 | **0.026** |

### Type of response and proximity (Hypothesis 4)

We next examined how the rate of response type to the gestures (response by visual or tactile, gesture, response by vocalization, response by activity change) was associated with the proximity (Table 8). There was a significant negative association between proximity and response by visual or tactile gesture (β = − 0.391, *p* = 0.012). In contrast, there was a significant positive association between the proximity and response by activity change (β = 0.603, *p* = 0.002).

**Table 8.** MRQAP regression models showing predictors of proximity (duration spent within 10 meters per hour spent in same party) by rate of response to the gesture between N = 12, 132 dyadic relationships of the chimpanzees. Significant P values are indicated in bold.

| Attribute category/ rate of gesture sequence per hour spent in close proximity | Standardized coefficient | Standard error | P |
| --- | --- | --- | --- |
| Age similarity | 0.198 | 3.887 | 0.026 |
| Sex similarity | -0.127 | 3.802 | 0.154 |
| Kinship similarity | 0.063 | 6.539 | 0.239 |
| Oestrous similarity | -0.004 | 4.093 | 0.479 |
| Response by visual or tactile gesture | -0.391 | 6.567 | **0.012** |
| Response by activity change | 0.603 | 1.746 | **0.002** |
| Response by vocalisation | -0.088 | 0.761 | 0.198 |

### Sequence network size and centrality in proximity network (Hypothesis 5)

Finally, we used node-level regressions to examine the association between gesture sequences (rapid and persistence), single gestures and individual position in the proximity network (centrality outdegree). Out degree refers to behaviours directed by the focal chimpanzee to conspecifics, whilst in degree refers to behaviours directed by conspecifics towards the focal chimpanzee. The network can vary between dyad A to B and B to A (e.g. proximity of Bwoba to Hawa can be different from proximity of Hawa to Bwoba), therefore in degree and out degree are calculated separately. All analyses controlled for the duration of time spent in proximity to oestrus females, time spent in proximity to kin, and the age and sex of the focal chimpanzee. We found that there was a positive association between the proximity outdegree and a persistence sequence indegree (β = 1.858, *p* = 0.015, Table 9). Thus individual chimpanzees with a higher rate of social behaviours directed at them also received a higher rate of persistence sequence directed at them.

**Table 9.** Node-level regression models predicting proximity out (overall durations of time spent in proximity within 10 meters per hour dyad spent in same party produced). Out degree refers to behaviours directed by the focal chimpanzee to conspecifics, whilst in degree refers to behaviours directed by conspecifics towards the focal chimpanzee. Based on 12 chimpanzees. Significant *p* values are indicated in bold.

| Attribute category/ Agreement in gesture repertoires | Standardized coefficient | *P* |
| --- | --- | --- |
| Reproductive state of female | -1.605 | **0.025** |
| Kinship | 0.359 | 0.250 |
| Sex/ age | -0.492 | 0.210 |
| Rapid sequence outdegree | -0.112 | 0.466 |
| Rapid sequence indegree | -0.046 | 0.471 |
| Single gesture outdegree | 0.255 | 0.431 |
| Single gesture indegree | -0.691 | 0.166 |
| Persistence sequence outdegree | -0.208 | 0.389 |
| Persistence sequence indegree | 1.858 | **0.015** |

## Discussion

An important aspect in understanding the evolution of complex sociality in humans is to understand the role of primate sequences of gestures in social bonding at the level of the dyad and the group. Primates produce single gestures (produced singly rather than in series), persistence sequences (series of gestures interspersed with periods of response waiting) and rapid sequences (series of gestures made in quick succession without periods of response waiting) (Hobaiter and Byrne 2011b). Recent theoretical accounts emphasize the role of gestures not purely as a means of information transfer (Seyfarth et al. 2010), but as a time-efficient mechanism of social bonding (Dunbar 2012). However, studies to date have examined the characteristics of gesture in isolation from the social system (Bard 2017; Bard et al. 2017; Byrne et al. 2017; Leavens et al. 2017). Thus, the mechanisms through which gesture sequences can be associated with more complex social systems remain unresolved. In this study, we used a sample of twelve wild chimpanzees to examine how the production of gesture sequences is related to patterns of sociality at both the individual and group levels. This extends previous research which has focused on the associated between type of gesture sequence and the response of the recipient. Overall the results demonstrate a significant association between patterns of proximity between pairs of chimpanzees and rates of different types of gestural communication.

Longer durations of proximity, per hour they spent within 10m, were associated with a higher rate of persistence sequence, but not a higher rate of single gesture or rapid sequences, supporting Hypothesis 1. Further, higher rates of intentional gestures (persistence, single gesture) were associated with response by activity change, whereas higher rate of emotional gestures (rapid sequence) were associated with response by vocalisation supporting Hypothesis 2. Finally, longer durations of proximity were associated with a higher rates of response present and response by means of activity change, supporting Hypotheses 3 and 4. These results suggest that one possible function of communication between individuals who spent a longer duration of time in social behaviors is to enable behavioral synchrony by influencing the behaviour of the recipient in goal directed and intentional way. Thus, one important dimension of complex social interactions is the degree of successful inter-individual adjustment between interactants, enabling them to coordinate joint activities such as travel or mutual grooming through intentional gesturing. Recent studies have provided evidence that gestural communication responded to by the recipient appears to be related to stronger social bonds than communication which has not been responded to (Schneider et al. 2017). Therefore one reason why individuals who spent longer durations of time in proximity use intentional gestures is because they can influence recipient flexibly to facilitate social interaction and achieve the communication goal (Roberts et al. 2014a).

In line with previous research in this area (Lehmann et al. 2016; Sapolsky et al. 1997; Silk et al. 2013; Silk et al. 2010b) we used proximity to measure the level of sociality of pairs of chimpanzees. This allowed for the association between sociality and rates of different types of gestural communication to be explored. However, different types of social behaviors may play a different roles in social cohesion in primates. For instance, the role of grooming in primate social relationships is well established (Dunbar 2010), but the role of other joint behaviours such as joint travel or joint feeding is less clear (Gruber and Zuberbühler 2013; King et al. 2011). Similarly, in humans cooperative contexts whereby actors co-regulate behaviour with one another to achieve common goal (e.g. joint travel) reflect stronger social bonding than other contexts (Pollet et al. 2013; Wolf et al. 2016). However, whether these different dimensions of sociality are differentially linked to gestural communication within dyads is unclear from this study and future work could examine specific instances of behaviour (e.g. grooming interactions, travel initiation) to explore the role of different types of gestural communication in coordinating this behaviour (Fedurek et al. 2015).

This interpersonal adjustment in social relationships may be particularly cognitively demanding and this this is especially the case in species where individuals have to manage a larger number of differentiated social relationships (Dunbar 2012; Freeberg et al. 2012). In particular, fission-fussion societies, and species of primates that live in large social groups, face cognitive challenges in maintaining a differentiated social network consisting of both stronger and weaker social ties (Silk et al. 2010a). Maintaining these different types of social bonds is necessary to preserve group cohesion (Henzi et al. 1997). In this study, we found that individual differences in sociality (centrality in the network) was associated with different patterns of gestural communication, supporting Hypothesis 5. Specifically centrality was positively associated with the rate of persistence sequence directed at the central individual, but not the rate of single gesture or rapid sequences directed at the central individual. This suggests that flexible use of persistence sequences is valuable to ensure the goal of communication is met (Roberts et al. 2014a) and intentional gestures play an important role in chimpanzee sociality.

The conclusions drawn in this study could be influenced by the uneven representation of different gestures within dataset. Previous studies which employed continuous observation of gestures have ranged between three (Hobaiter et al. 2017) to five hours (Wilke et al. 2017) of observation of each focal individual during study period. In the current study, we observed 12 focal subjects from a single study group for a mean duration of 12 hours per each focal chimpanzee, ranging between 8.3 hours to 18.63 hours (taking into account the video data collected in parallel with the socio-ecological samples during the last data collection season). However, the sampling of focal individuals was uneven and single gestures and sequences vary in their occurrence rates. For instance, in this study, there were 160 sequences of different types whereas there were 385 single gestures. Similarly, gesture types were not distributed evenly across categories, as a majority of gesture types were confined to most common occurrence categories. Thus whilst the results are broadly in line linking gestural communication with sociality and coordination of behavior in primates (Byrne et al. 2017; Leavens et al. 2005; Roberts et al. 2014b), further research is needed to explore how gestural communication is associated with sociality in other chimpanzee communities and other primate species. This further research could focus on compiling a dataset whereby gesture sequences and gesture types would be represented more equally. Further, whilst we explored associations between sociality and gestural communication, we could not demonstrate a causal relationship between gestural communication and a longer durations of proximity between pairs of chimpanzees. Research examining how specific types of gestural communication are associated with the durations of specific instances of social behavior would be needed to establish such a causal relationship. Many gestures are produced in the context of grooming (Byrne et al. 2017; Roberts et al. 2012a) and one promising area for future research would be to examine whether specific types of gestures given in grooming contexts are associated with longer grooming bouts or reduced probability of defecting to an alternative grooming partner (Fedurek et al. 2015; Kaburu and Newton-Fisher 2016)

The predictability of conspecifics’ behaviour is a major modulator of stress in group living animals (Seyfarth and Cheney 2013) and greater use of intentional gestures may reduce this stress by increasing the likelihood of the recipient responding appropriately to the gesture. This is especially important as gestural communication can be used in both affiliative and agonistic contexts in close proximity and thus intentional gestures may lead to greater coordination between the pair of chimpanzees. Previous research has focused on how intentionality in gestural communication is related to the recipients’ response and comprehension of signaling, both in relation to human and conspecific recipients. Whilst this research has detailed the extent to which chimpanzees can flexibility adjust their communication, and explored how sensitive these adjustments are to different aspects of the recipients response, it has not demonstrated how this flexibility in communication helps chimpanzees meet the key adaptive challenges faced by group living animals – maintaining a differentiated set of stable, long-term social relationships and responding appropriately to others (Dunbar and Shultz 2007a). If the key driving force of brain evolution in both primates and hominins has been evolution of complex social relationships rather than ecological factors (Dunbar and Shultz 2007b), the cognitive skills underpinning flexibility in communication should enable primates to meet these social challenges. The current results suggest that intentional gestural communication may enable greater levels of behavioural coordination when interacting at close proximity and thus longer durations of proximity and affiliative activities such as grooming.

To conclude, the ability to accurately coordinate social behavior through gestural signals with conspecifics is a key aspect of successful group living (Seyfarth and Cheney 2013). The findings of this study demonstrated that flexibility in gestural communication is associated with sociality and may help chimpanzees meet the challenges of group living, with persistence in particular being associated with longer durations of proximity. Individual variation in the strength of social bonds in primates is strongly linked to fitness outcomes (Silk 2007) and our results suggest that flexibility in gestural communication may play an important role in explaining some of this individual variation in social relationships.

## References

Bakeman R, Gottman JM (1997) Observing interaction: An introduction to sequential analysis. Cambridge university press,

Bard KA (1992) Intentional Behavior and Intentional Communication in Young Freeberg Ranging Orangutans Child Development 63: 1186–1197

Bard KA (2017) Dyadic interactions, attachment and the presence of triadic interactions in chimpanzees and humans Infant Behavior and Development 48: 13–19

Bard KA, Dunbar S, Maguire-Herring V, Veira Y, Hayes KG, McDonald K (2014) Gestures and social-emotional communicative development in chimpanzee infants American Journal of Primatology 76: 14–29

Bard KA, Maguire-Herring V, Tomonaga M, Matsuzawa T (2017) The gesture ‘Touch’: Does meaning-making develop in chimpanzees’ use of a very flexible gesture? Animal cognition:1–16

Bates E, Benigni L, Bretherton I, Camaioni L, Volterra V (1979) The emergence of symbols. Academic Press, New York

Borgatti SP, Everett MG, Freeman LC (2014) Ucinet. In: Alhajj R, Rokne J (eds) Encyclopedia of Social Network Analysis and Mining. Springer-Verlag New York, pp 2261–2267

Borgatti SP, Everett MG, Johnson JC (2013) Analyzing Social Networks. SAGE Publications Limited,

Byrne RW, Cartmill E, Genty E, Graham KE, Hobaiter C, Tanner J (2017) Great ape gestures: intentional communication with a rich set of innate signals Animal Cognition:1–15 doi:10.1007/s10071-017-1096-4

Cartmill E, Byrne R (2007a) Orangutans modify their gestural signaling according to their audience’s comprehension Current Biology 17: 1345–1348

Cartmill EA, Byrne RW (2007b) Orangutans modify their gestural signaling according to their audience’s comprehension Current Biology 17:1345–1348 doi:10.1016/j.cub.2007.06.069

Cartmill EA, Byrne RW (2010) Semantics of primate gestures: intentional meanings of orangutan gestures Animal Cognition 13: 793–804

Dekker D, Krackhardt D, Snijders TA (2007) Sensitivity of MRQAP tests to collinearity and autocorrelation conditions Psychometrika 72: 563–581

Dunbar R (2012) Bridging the bonding gap: The transition from primates to humans Philosophical Transactions of the Royal Society B: Biological Sciences 367: 1837–1846

Dunbar RI, Shultz S (2007a) Evolution in the social brain Science 317: 1344–1347

Dunbar RI, Shultz S (2007b) Understanding primate brain evolution Philosophical Transactions of the Royal Society of London B: Biological Sciences 362: 649–658

Dunbar RIM (1993) Coevolution of neocortical size, group size and language in humans Behavioral and Brain Sciences 16: 681–694

Dunbar RIM (1998) The social brain hypothesis Evolutionary Anthropology 6: 178–190

Dunbar RIM (2010) The social role of touch in humans and primates: Behavioural function and neurobiological mechanisms Neuroscience & Biobehavioral Reviews 34: 260–268

Fedurek P, Slocombe KE, Hartel JA, Zuberbühler K (2015) Chimpanzee lip-smacking facilitates cooperative behaviour Scientific reports 5: 13460

Forrester SG (2008) A multidimensional approach to investigations of behaviour: Revealing structure in animal communication signals Animal Behaviour 76: 1749–1760

Freeberg TM, Dunbar RI, Ord TJ (2012) Social complexity as a proximate and ultimate factor in communicative complexity Philosophical Transactions of the Royal Society B: Biological Sciences 367: 1785–1801

Fröhlich M, Wittig RM, Pika S (2016) Play-solicitation gestures in chimpanzees in the wild: flexible adjustment to social circumstances and individual matrices Royal Society Open Science 3: 160278

Genty E, Breuer T, Hobaiter C, Byrne RW (2009) Gestural communication of the gorilla (*Gorilla gorilla*): Repertoire, intentionality and possible origins Animal Cognition 12:527–546 doi:10.1007/s10071-009-0213-4

Genty E, Byrne RW (2009) Why do gorillas make sequences of gestures? Animal Cognition 13: 287–301

Gillespie-Lynch K, Greenfield PM, Feng Y, Savage-Rumbaugh S, Lyn H (2013) A cross-species study of gesture and its role in symbolic development: implications for the gestural theory of language evolution Front Psychol 4:10.3389

Gómez JC (1996) Ostensive behaviour in great apes: The role of eye contact. In: Russon AE, Bard KA, Parker ST (eds) Reaching into Thought. The Minds of the Great Apes. Cambridge University Press, Cambridge,

Goodall J (1986) The Chimpanzees of Gombe: Patterns of Behaviour. Harward University Press, Cambridge, Massachusetts

Gruber T, Zuberbühler K (2013) Vocal recruitment for joint travel in wild chimpanzees PLoS One 8:e76073

Halina M, Rossano F, Tomasello M (2013) The ontogenetic ritualization of bonobo gestures Animal cognition 16: 653–666

Henzi S, Lycett J, Piper S (1997) Fission and troop size in a mountain baboon population Animal Behaviour 53: 525–535

Hewes GW (1973) Primate communication and the gestural origin of language Current Anthropology 14: 5–24

Hill RA, Bentley RA, Dunbar RI (2008) Network scaling reveals consistent fractal pattern in hierarchical mammalian societies Biology letters 4: 748–751

Hobaiter C, Byrne RW, Zuberbühler K (2017) Wild chimpanzees’ use of single and combined vocal and gestural signals Behavioral Ecology and Sociobiology 71: 96

Hobaiter K, Byrne R (2011a) The gestural repertoire of the wild chimpanzee Animal Cognition 14:745–767 doi:10.1007/s10071-011-0409-2

Hobaiter K, Byrne R (2011b) Serial gesturing by wild chimpanzees: Its nature and function for communication Animal Cognition 14:827–838 doi:10.1007/s10071-011-0416-3

Kaburu SS, Newton-Fisher NE (2016) Bystanders, parcelling, and an absence of trust in the grooming interactions of wild male chimpanzees Scientific reports 6

King AJ, Clark FE, Cowlishaw G (2011) The dining etiquette of desert baboons: the roles of social bonds, kinship, and dominance in co-feeding networks American Journal of Primatology 73: 768–774

Langergraber K, Mitani J, Vigilant L (2009) Kinship and social bonds in female chimpanzees (Pan troglodytes) American Journal of Primatology 71: 840–851

Leavens DA, Bard KA, Hopkins WD (2017) The mismeasure of ape social cognition Animal Cognition:1–18

Leavens DA, Russell JL, Hopkins WD (2005) Intentionality as measured in the persistence and elaboration of communication by chimpanzees (Pan troglodytes) Child Development 76: 291–306

Lehmann J, Majolo B, McFarland R (2016) The effects of social network position on the survival of wild Barbary macaques, Macaca sylvanus Behavioral Ecology 27: 20–28

Liebal K, Call J, Tomasello M (2004) Use of gesture sequences in chimpanzees American Journal of Primatology 64: 377–396

Maestripieri D (2005) Gestural communication in three species of macaques (Macaca mulatta, M. nemestrina, M. arctoides): Use of signals in relation to dominance and social context Gesture 5: 55–71

McCarthy MS, Jensvold MLA, Fouts DH (2012) Use of gesture sequences in captive chimpanzee (Pan troglodytes) play Animal Cognition:1–11

Mitani JC (2009) Male chimpanzees form enduring and equitable social bonds Animal Behaviour 77: 633–640

Mitani JC, Watts DP, Pepper JW, Merriwether DA (2002) Demographic and social constraints on male chimpanzee behaviour Animal Behaviour 64: 727–737

Moore R (2016) Meaning and ostension in great ape gestural communication Animal cognition 19: 223–231

Owren M, Rendall D (2001) Sound on the reboud: Bringing form and function back to the forefront in understanding nonhuman primate vocal signaling Evolutionary Anthropology 10: 58–71

Pika S, Liebal K, Tomasello M (2005) Gestural communication in subadult bonobos (*Pan paniscus*) : Repertoire and use American Journal of Primatology 65: 39–61

Pollet TV, Roberts SG, Dunbar RI (2013) Going that extra mile: individuals travel further to maintain face-to-face contact with highly related kin than with less related kin PloS one 8:e53929

Pollick AS, de Waal FBM (2007) Ape gestures and language evolution Proceedings of the National Academy of Sciences of the United States of America 104: 8184–8189

Reynolds V (2005) The chimpanzees of the Budongo Forest: ecology, behaviour, and conservation. Oxford University Press, Oxford

Roberts AI, Roberts SGB (2016a) Wild chimpanzees modify modality of gestures according to the strength of social bonds and personal network size Scientific Reports 6 doi:10.1038/srep33864

Roberts AI, Roberts SGB, Vick S-J (2014a) The repertoire and intentionality of gestural communication in wild chimpanzees Animal Cognition 17:317 - 336 doi:10.1007/s10071-013-0664-5

Roberts AI, Vick S-J, Buchanan-Smith H (2012a) Usage and comprehension of manual gestures in wild chimpanzees Animal Behaviour 84:459–470 doi:10.1016/j.anbehav.2012.05.022

Roberts AI, Vick S-J, Buchanan-Smith H (2013) Communicative intentions in wild chimpanzees: Persistence and elaboration in gestural signalling Animal Cognition 16:187–196 doi: 10.1007/s10071-012-0563-1

Roberts AI, Vick S-J, Roberts SGB, Buchanan-Smith HM, Zuberbühler K (2012b) A structure-based repertoire of manual gestures in wild chimpanzees: Statistical analyses of a graded communication system Evolution and Human Behavior 33:578–589 doi:10.1016/j.evolhumbehav.2012.05.006

Roberts AI, Vick S-J, Roberts SGB, Menzel CR (2014b) Chimpanzees modify intentional gestures to coordinate a search for hidden food Nature Communications 5 3088 doi:10.1038.ncomms4088

Roberts SGB, Roberts AI (2016b) Social brain hypothesis, vocal and gesture networks of wild chimpanzees Frontiers in Psychology 7 doi:10.3389/fpsyg.2016.01756

Sapolsky RM, Alberts SC, Altmann J (1997) Hypercortisolism associated with social subordinance or social isolation among wild baboons Archives of General Psychiatry 54: 1137–1143

Schneider C, Call J, Liebal K (2012) Onset and early use of gestural communication in nonhuman great apes American journal of primatology 74: 102–113

Schneider C, Liebal K, Call J (2017) “Giving” and “responding” differences in gestural communication between nonhuman great ape mothers and infants Developmental Psychobiology 59: 303–313

Scott-Phillips TC (2015a) Meaning in animal and human communication Animal cognition 18: 801–805

Scott-Phillips TC (2015b) Nonhuman primate communication, pragmatics, and the origins of language Current Anthropology 56: 56–80

Scott N (2013) Gesture Use by Chimpanzees (Pan troglodytes): Differences Between Sexes in Inter- and Intra-Sexual Interactions American Journal of Primatology 75: 555–567

Seyfarth RM, Cheney DL (2013) Affiliation, empathy, and the origins of theory of mind Proceedings of the National Academy of Sciences 110: 10349–10356

Seyfarth RM, Cheney DL, Bergman T, Fischer J, Zuberbuhler K, Hammerschmidt K (2010) The central importance of information in studies of animal communication Animal Behaviour 80:3–8 doi:DOI 10.1016/j.anbehav.2010.04.012

Silk J, Cheney D, Seyfarth R (2013) A practical guide to the study of social relationships Evolutionary Anthropology: Issues, News, and Reviews 22: 213–225

Silk JB (2007) Social components of fitness in primate groups Science 317: 1347–1351

Silk JB et al. (2010a) Female chacma baboons form strong, equitable, and enduring social bonds Behavioral Ecology and Sociobiology 64: 1733–1747

Silk JB et al.(2010b) Strong and consistent social bonds enhance the longevity of female baboons Current Biology 20: 1359–1361

Spoor JR, Kelly JR (2004) The evolutionary significance of affect in groups: Communication and group bonding Group processes & intergroup relations 7: 398–412

Taglialatela JP et al.(2015) Multimodal communication in chimpanzees American journal of primatology 77: 1143–1148

Tanner JE (2004) Gestural phrases and gestural exchanges by a pair of zoo-living lowland gorillas Gesture 4: 1–24

Tanner JE, Perlman M (2016) Moving beyond ‘meaning’: Gorillas combine gestures into sequences for creative display Language & Communication:1 - 17 doi:10.1016/j.langcom.2016.10.006

Tempelmann S, Liebal K (2012) Spontaneous use of gesture sequences in orangutans Developments in Primate Gesture Research 6: 73

Tomasello M, Call J, Nagell K, Olguin R, Carpenter M (1984) The learning and use of gestural signals by young chimpanzees: A trans-generational study. Primates 37: 137–154.

Tomasello M, George BL, Kruger AC, Jeffrey M, Evans FA (1985) The development of gestural communication in young chimpanzees Journal of Human Evolution 14: 175–186

Tomasello M, Zuberbühler K (2002) Primate vocal and gestural communication. In: Bekoff M, Allen CS, Burghardt G (eds) The cognitive animal: empirical and theoretical perspectives on animal cognition. MIT Press, Cambridge,

Townsend SW et al.(2016) Exorcising Grice’s ghost: an empirical approach to studying intentional communication in animals Biological Reviews

Wilke C, Kavanagh E, Donnellan E, Waller BM, Machanda ZP, Slocombe KE (2017) Production of and responses to unimodal and multimodal signals in wild chimpanzees, Pan troglodytes schweinfurthii Animal Behaviour 123: 305–316

Wolf W, Launay J, Dunbar RI (2016) Joint attention, shared goals, and social bonding British Journal of Psychology 107: 322–337

